# Stochastic Evolutionary Control in Heterogeneous Populations

**DOI:** 10.64898/2026.03.12.711493

**Authors:** Peng Chen, Jonathan Asher Pachter, Jacob G. Scott, Michael Hinczewski

**Affiliations:** Department of Physics, Case Western Reserve University, Cleveland, OH 44106; Genomic Sciences and Systems Biology, Cleveland Clinic Research, Cleveland, OH 44106; School of Medicine, Case Western Reserve University, Cleveland, OH 44106

**Author notes:** These authors contributed equally to this work (Peng Chen, Jonathan Asher Pachter).

**Keywords:** evolutionary control, Wright–Fisher dynamics, Markov Decision Process, drug resistance

## Abstract

Therapeutic resistance, which poses a central challenge in cancer and infectious disease treatment, arises from the evolutionary dynamics of heterogeneous populations as a natural consequence of the evolutionary tendency towards higher fitness. Here we introduce SHEPHERD (Stochastic Heterogeneity–informed Evolutionary Policy Hampering the Expansion of Resistance to Drugs), which integrates Wright–Fisher (WF) population genetics modeling with Markov Decision Processes (MDPs) to design optimal adaptive drug policies which reduce the fitness of evolving disease, mitigating resistance. While MDPs have been applied to evolutionary control under the strong-selection, weak-mutation (SSWM) assumption—where populations are effectively homogeneous—SHEPHERD operates beyond this regime, effectively capturing the full stochastic dynamics of genetically heterogeneous populations experiencing mutation and selection of a wide array of strengths. Using synthetic multi-drug fitness landscapes, we show the benefit of the optimized SHEPHERD protocol for *in silico* disease systems with three, four, and eight genotypes: the long-term mean fitness of the disease is reduced compared to any single-drug or two-drug periodic regimen. Although these examples specifically model drug resistance, we discuss the potential broad applicability of our approach to calculating strategies for controlling a wide class of evolving populations. We further analyze the sensitivity of SHEPHERD strategies to different levels of genotypic frequency discretization and temporal resolution, demonstrating robustness once moderate resolution is achieved, but strong dependence on the timing of drug updates. Although current results rely on synthetic landscapes due to the lack of empirical multi-genotype, multi-drug measurements outside the SSWM regime, our study establishes a foundation for applying MDP optimization to broad classes of evolutionary dynamics, highlighting both the opportunities and the computational challenges of controlling evolution.

## 1. Introduction

Understanding how evolutionary processes are influenced by shifting environmental conditions is a fundamental scientific challenge with wide-ranging societal implications. Whether in the context of climate change^1,2^, conservation of endangered species^3^, or agriculture^4^, changing external conditions impact evolution in complex and stochastic ways. Understanding this complexity would beget the ability to predict, and ultimately intervene with conscience and purpose to steer these complex systems toward desired objectives^5^. This ability to enact evolutionary control could be especially beneficial in the realm of human disease.

Certain diseases can evolve resistance to treatment—a phenomenon paradoxically induced by the very treatment that sought to eradicate the disease. This is a major public health concern and an area of active research, from cancer^6–8^to microbes^9–12^. One avenue for combating this issue is harnessing the evolution of the disease to our own ends. In the process of developing resistance to one drug, a disease may become particularly susceptible to a second drug, a phenomenon known as “collateral sensitivity”^13–16^; this suggests the ability to steer the evolution of a disease by carefully choosing sequences of drugs to administer^17–21^. Indeed, this has inspired theoretical and experimental work exploring how to find optimal treatment protocols in the presence of collateral sensitivity^22–25^.

Mathematical modeling has, in general, played a central role in studying progression of cancer and infectious diseases and their response to intervention^26–29^. There is a rich history of research into therapeutic interventions optimally controlling evolving diseases^5,30–33^including cancer^34–37^and bacterial infections^38,39^, as well as viruses such as HIV^40–44^. Some proposed drug protocols are adaptive, maximizing potential success by responding to the state of the disease at a given time^45–50^.

While it is by no means universal, underlying models of disease evolution are often deterministic, captured in ordinary differential equations^51^, whereas evolution is a fundamentally stochastic process^52^. An apt framework for describing adaptive control in such stochastic systems is Markov Decision Processes (MDPs). As the name implies, MDPs combine a stochastic Markov process with an element of control: at each time step, a choice is made which influences the probabilities for the next transition of the stochastic system^53^. MDPs are incredibly versatile—they underlie reinforcement learning, they have applications in robotics, and they have been already used for years in medical contexts^54,55^such as mitigating side effects of cancer treatment^56,57^, as well as successfully modeling evolving diseases exposed to different drugs^23,25^.

A recent paper from our group^58^ specifically employed MDPs to learn about optimal drug protocols for treating homogeneous populations of disease cells, i.e. all cells in the population were assumed to have the same genotype at any given time, though that genotype can change due to a mutation that fixes in the population – this is an approximation in the regime of strong selection and weak mutation (SSWM). While this work was an important step that provides insight to certain situations, cancer in particular is known to display significant genetic heterogeneity^8,59–61^. Similarly, heterogeneity is not always included in the mathematical studies of drug resistance in microbial disease^51^. Thus, we desire a mathematical model that incorporates genetic heterogeneity—namely, we use the Wright-Fisher (WF) model^62,63^, a stalwart of population genetics, which allows for phenomena such as clonal interference and competition among genotypes, and which has been employed successfully to model evolving diseases including cancer^32,64–67^. A previous study did employ WF modeling in optimal control of disease^30^, though it still primarily considered strong selection and was limited to two genotypes and two control actions at a time – constraints we move beyond in this work.

In summary, we seek to build upon the wealth of existing work, bridging gaps and combining ingredients in a novel way. To that end, this paper introduces the SHEPHERD protocol (Stochastic Heterogeneity–informed Evolutionary Policy Hampering the Expansion of Resistance to Drugs), which computes optimal, adaptive drug protocols for evolving, genetically heterogeneous populations—combining the complexity of WF modeling with the power of the MDP framework, and the evolutionary steerability afforded by cycling between multiple, potentially collaterally sensitive drugs. Additionally, the SHEPHERD protocol contains novelty in its approach to computing the optimal MDP, employing mathematical approximations that simplify and speed up calculations without overly sacrificing performance, as detailed below.

In Section 2, we outline the mathematical foundations of WF modeling and MDPs as they apply to drug protocols, and describe how these elements integrate into the SHEPHERD protocol for computing optimal drug protocols, including outlining the mathematical approximations used to enhance tractability of calculations. Subsequently, in Section 3, we present results from numerical calculations exhibiting the power of SHEPHERD, with metrics of performance such as the clear improvement over both single-drug and two-drug switching protocols. Finally, in Section 4, we discuss the implications of our results. In particular, we note that, while this work was motivated by drug resistance in cancer and other diseases, the SHEPHERD approach is much more general: it can compute strategies to make optimal use of any external controlling factors for achieving arbitrary goals in WF evolving populations.

## 2. Mathematical Elements of the SHEPHERD Protocol

### 2.1 A Special Coarse-Graining of Wright-Fisher Dynamics

As mentioned above, we employ the Wright-Fisher (WF) model of evolutionary dynamics (see **Box 1** for review). In the following subsection, we describe three steps we took to improve our ability to simulate WF dynamics and compute optimal policies on said dynamics: a diffusion approximation, followed by a coordinate transformation that diagonalizes the diffusion equation, followed by a lattice discretization. While we both start and end with a discrete description of the dynamics, the final result is greatly coarse-grained and simplified with respect to the original description of the process; this allows us to calculate optimal control policies for larger population sizes using less computing power and time (since we have reduced dimensionality) but without sacrificing the ability to generate successful control policies, which we explicitly demonstrate by testing the control policies calculated under approximation on the original WF dynamics, a non-trivial validation of our method.

**Box 1 | Wright-Fisher Model: Stochastic Heterogeneous Evolving Populations**

We briefly review the Wright-Fisher (WF) model of evolution^62,63^, upon which the methods of this paper are based. Consider a finite population of *N* asexually reproducing individuals, comprising *M* distinct genotypes; this is a very general premise, capable of describing any asexually reproducing finite population, but we will consider for now populations of evolving disease units—cancer cells in tumors or infections of bacterial cells or even viruses, i.e. collections of asexually reproducing units causing harm to humans. One simplifying assumption of WF modeling is constant population size (i.e. a population at the carrying capacity of its environment), though in some cases varying population models can be mapped onto a WF process with constant effective population size^68,69^.

Let ***x***_*t*_ = (*x*_*t*,0_, …, *x*_*t,M*−1_) denote the genotype frequencies in the population at generation *t*, with 0 ≤ *x*_*t,I*_ ≤ 1 and ∑_*i*_ *x*_*t,i*_ = 1. In the classical WF process, the frequencies ***x***_*t*_ are updated in discrete generations through three steps:

1. *Selection*. Individuals reproduce in proportion to their reproductive fitness, where the fitness vector ***f*** = ( *f*_0_, …, *f*_*M*−1_) denotes the fitness values *f*_*i*_ for each genotype *i*, so the mean fitness at time *t* for a particular genetic composition ***x***_*t*_ is 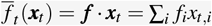. Mathematically, this step involves the transformation of frequencies *x*_*t,i*_ to fitness-biased frequencies 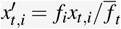 Different genotypes of disease exhibit different fitness when exposed to different drugs, quantifying their resistance or sensitivity to the treatment. We model therapeutic interventions by assigning to each drug a fitness vector 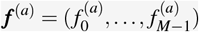, which defines the corresponding fitness landscape: genotype *i* has reproductive success 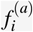 under drug *a*. (Note that we can also consider control actions as representing the same drug administered at distinct doses, or even no drug at all.)
2. *Mutation*. Offspring genotypes may mutate according to a stochastic mutation matrix 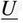, where the element *U*_*ji*_ is the probability for genotype *i* to mutate to genotype *j* for *i ≠ j*, and the diagonal terms are fixed by the columns summing to one. Mathematically, this means the fitness-biased frequencies 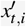 now become the mutated and fitness-biased frequencies 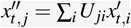. In this work, we often model genotypes as bit strings, with zeroes corresponding to the wild-type gene at that locus and ones corresponding to mutations at that locus. In that case, we typically use a Hamming-1 mutation graph, where each genotype mutates to its single-bit neighbors with probability *µ* per generation^70,71^, i.e. mutations in genes of interest only occur at one locus at most per reproductive event. However, the approach is applicable to any mutation matrix representing any mutation graph topology; for example, in the three-genotype example discussed below, we consider symmetric mutation between all genotypes.
3. *Genetic drift*. From the post-selection, post-mutation distribution, the next generation of size *N* is sampled multinomially, introducing stochastic fluctuations due to finite population size. WF is thus a model of non-overlapping generations. (This formulation also assumes the population is well-mixed, i.e. there is no spatial or physical interaction structure to the population affecting reproduction.) Mathematically, putting it all together, this means that the probability to transition from frequencies ***x***_*t*_ at time *t* to frequencies ***x***_*t*+1_ at time *t* + 1 is

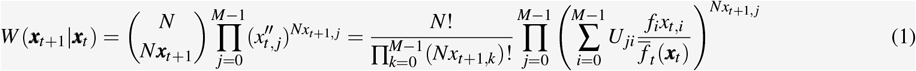

where the only possible new frequencies are those that still obey the positivity and normalization conditions 0 ≤ x_t+1,i_ ≤ 1 and ∑_i_ x_t+1,i_ = 1.

Together, these steps define the WF stochastic model of (drug-modulated) evolution, as depicted schematically in **Figure 1A** and **Figure 1D**.

**Figure 1.**
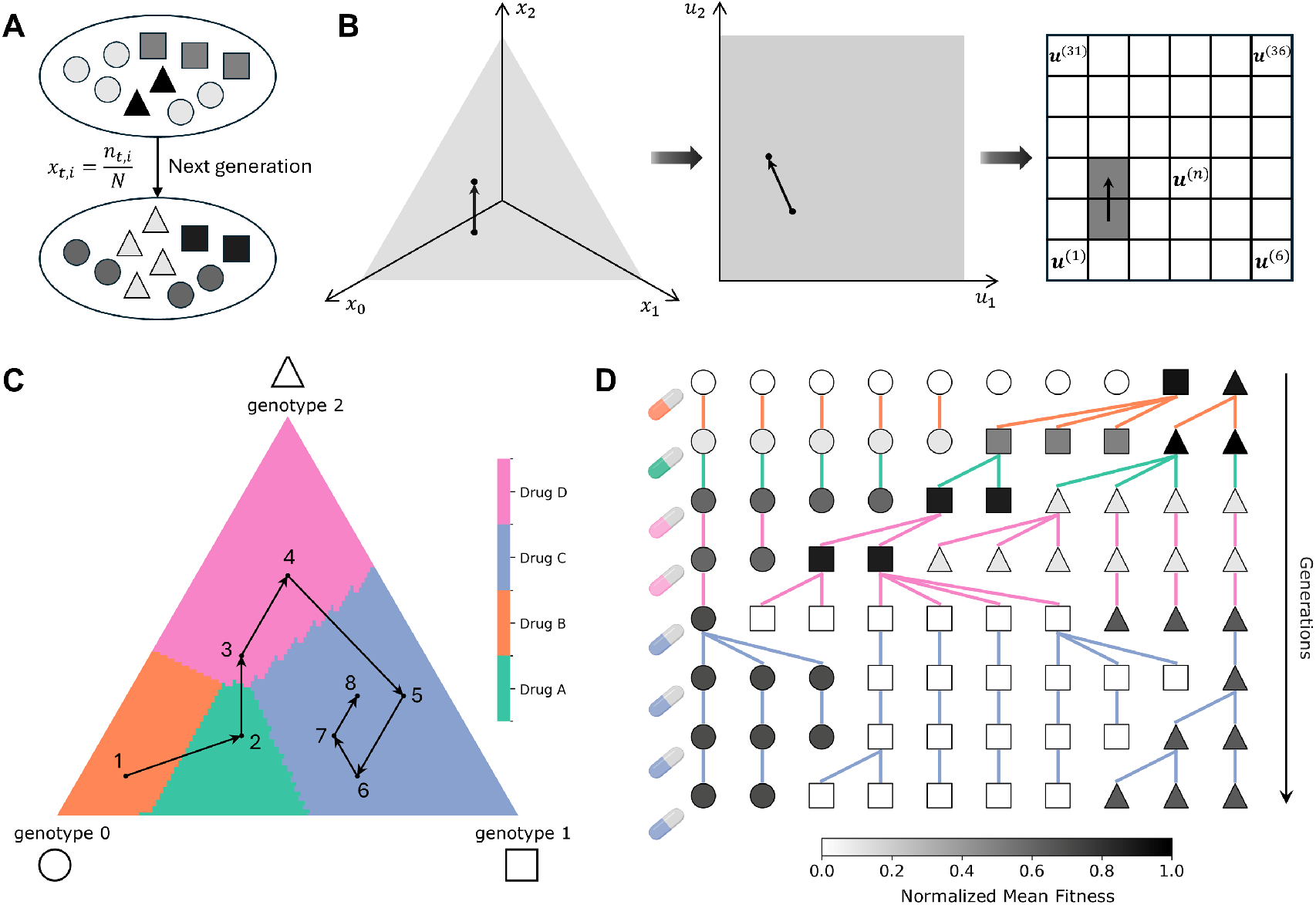
Schematic of the SHEPHERD protocol for evolutionary control. **A**. Wright–Fisher (WF) population dynamics illustrating the stochastic sampling of genotypes between generations. Each genotype is represented by a distinct shape, and its frequency is defined as *x*_*t,i*_ = *n*_*t,i*_*/N*, where *n*_*t,i*_ is the number of individuals of genotype *i* and *N* is the total population size. **B**. The two generations shown in panel A correspond to two points in genotype-frequency space (*x*_0_, *x*_1_, *x*_2_), which are transformed from the simplex to a square domain (*u*_1_, *u*_2_) and discretized into a finite grid for computation within the Markov Decision Process (MDP) framework. **C**. Optimal MDP policy defined over the genotype-frequency simplex, where each region is colored by the drug choice that minimizes long-term population fitness. The black trajectory (points 1–8) shows an example evolutionary path through genotype space under the policy. **D**. WF dynamics under the optimal MDP policy, where the applied drug may change in each generation based on the current genotype frequencies. The eight generations correspond to the trajectory shown in panel C. Each generation (row) contains ten cells with genotype indicated by shape, lines between generations denote evolutionary transitions under the influence of drugs indicated by color, and the grayscale shading of cells indicates normalized mean fitness—the policy steers the population’s evolution toward lower fitness.

#### 2.1.1 Diffusion Approximation

When population size *N* is large and mutation and selection are not too strong (see below for quantification), the WF dynamics are well-approximated by a diffusion process described by the following Fokker–Planck equation for the probability density *p*(***x***, *t*) of genotype frequencies ***x*** at time *t* ^72,73^:

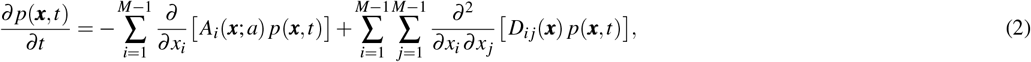

where ***A***(***x***; *a*) is the vector containing the deterministic part of the dynamics under the influence of drug *a* and 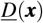 is the diffusion matrix, with time measured in units of generations of the underlying WF dynamics.

Mutation is encapsulated by a matrix 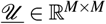, where 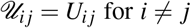 is the mutation rate from genotype *j* into genotype *i* (equal to the mutation probability *U*_*i j*_ per unit time of one generation), and 𝒰_*ii*_ is the negative sum of the rates ofmutation out of genotype *i* to all other genotypes (thus the columns sum to zero)^74^.

The deterministic and diffusion parts for the Fokker-Planck equation can then be written as ^72,73^

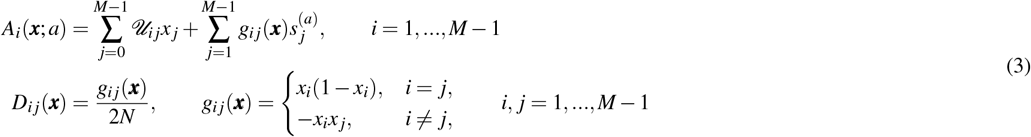

Where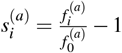are the relative selection coefficients, defined in terms of the absolute fitness 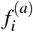of genotype *i* under drug *a* relative to the wild type 0 under the same drug. A positive value for 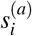 means type *i* is fitter than the wild type, while a negative value means it is less fit. Properly, for the Fokker-Planck equation to be a valid approximation of WF dynamics, all of the selection coefficients and mutation rates must satisfy must satisfy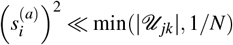, and all of the mutation rates must similarly satisfy (*𝒰*_*I j*_)^2^*≪*1*/N*; this ensures that expected changes in genotype frequencies are small enough to be approximately continuous for time measured in generations, and it also ensures that the diffusion term in the Fokker-Planck equation is independent of mutation and selection, merely representing the stochasticity arising from the multinomial sampling in a WF generational step.

To contrast with previous work employing very strong selection, our initial examples operate under very weak selection, in which selection coefficients satisfy 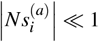for all genotypes *i* and drugs *a*; this regime corresponds to the significantly heterogeneous populations that we set out to model. However, we also show later that SHEPHERD successfully computes optimal control protocols in widely varying regimes of mutation and selection magnitude.

#### 2.1.2 Coordinate Transformation

The Fokker-Planck dynamics can be greatly simplified through an auspicious coordinate transformation^75^. We transform genotype frequencies ***x*** into new coordinates ***u*** that map the frequency simplex onto the unit hypercube [0, 1]^*M*−1^, which eases calculations. For *i* = 1, …, *M* − 1, we define

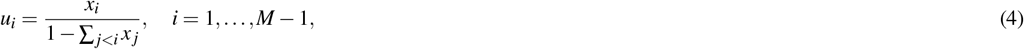

with the inverse transformation

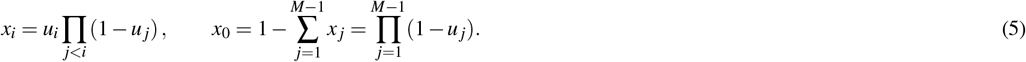

The Fokker-Planck equation in ***u***-space becomes^75^:

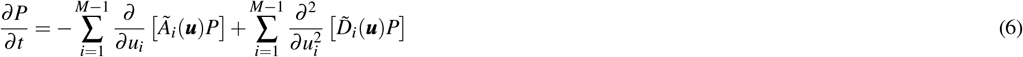

so that no mixed second–order derivatives appear; this diagonalization of the Fokker-Planck equation, combined with the move from simplex geometry to hypercube geometry, are the great benefits of the coordinate transformation (simplexes are, ironically, much less simple than hypercubes).

Defining *S*_*i*_(***u***) *≡* ∏ _*j<i*_(1 −*u*_*j*_) so that *x*_*i*_ = *u*_*i*_*S*_*i*_ for *i* ≥ 1 and *x*_0_ = *S*_*M*_, the diagonal diffusion term in the new coordinates is:

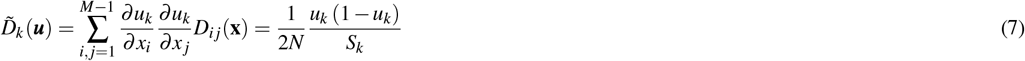

A derivation of the full form of the slightly more complicated deterministic part *Ã*_*k*_(***u***) in the new coordinates can be found in Supplementary Information II.

#### 2.1.3 Lattice Discretization

In the end, we still need to provide a discrete state space to numerically calculate the optimal control policy in the MDP framework, so we discretize the continuous Fokker–Planck dynamics by partitioning the unit hypercube [0, 1]^*M*−1^ into an *L×… ×L* = *L*^*M*−1^ lattice with spacing *h* along each axis. Lattice sites are indexed by the multi–index **i** = (*i*_1_, …, *i*_*M*−1_) with 1 ≤ *i*_*k*_ ≤ *L* for *k* = 1, …, *M* − 1, and mapped to a single index

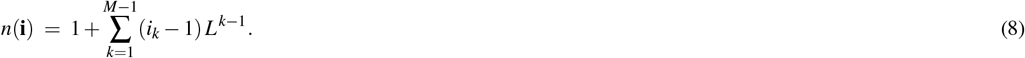

The corresponding cell–center coordinates are

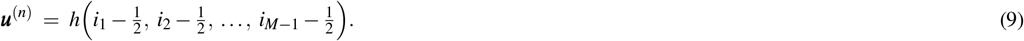

On this discretized state space, we construct a transition rate matrix 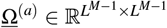using centered finite differences and nearest-neighbor jumps, which retrieves the Fokker-Planck dynamics in the limit as lattice spacing goes to zero (see Supplementary Information III for a detailed derivation). Specifically, for a forward jump along axis *k* we set

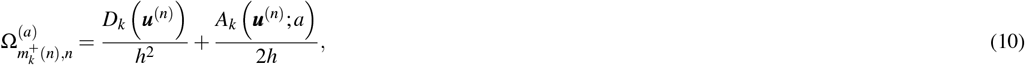

and for the corresponding backward jump

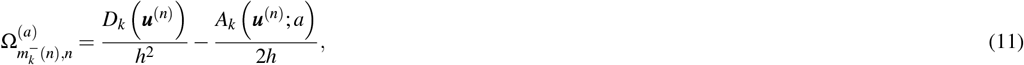

for each *k* = 1, …, *M*− 1, where 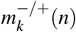labels the lattice position that is the same as the starting lattice position *n* except the *k*^th^ index *i*_*k*_ has been lowered/raised to *i*_*k*_ − */* + 1. All other off-diagonal entries are zero. The diagonal is chosen to conserve probability:

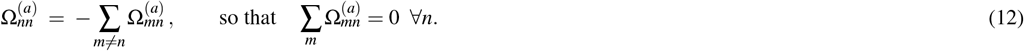

At the domain boundaries, we impose reflecting (no-flux) conditions by omitting jumps that would leave the domain (i.e., when *i*_*k*_ = 1 for a backward step or *i*_*k*_ = *L* for a forward step); the diagonal is still defined by the outgoing-rate sum above.

The transition–rate matrix 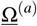specifies transition rates, i.e. transition probabilities per unit time of the Fokker-Planck dynamics. For calculating the optimal control policy in the MDP framework, we need transition probabilities for finite time steps Δ*t* corresponding to how long evolution occurs before a new action is chosen (in our case, when the drug is changed). The finite-time transition probability matrix is obtained by exponentiation of the transition rate matrix:

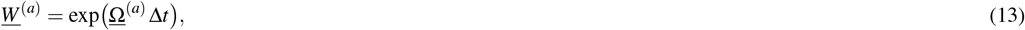

We typically use Δ*t* = 1, corresponding to one generation of the underlying WF process, but the formulation allows for arbitrary Δ*t* > 0, and in fact we later discuss the effects of varying Δ*t* in section 3.4.

Lastly, we note that these transition matrices can be very large, which is of course computationally costly and can impede some computations, especially for larger genotype spaces and higher lattice resolution. For *M* genotypes, there are *M* − 1 independent genotype frequencies; if these are discretized into *L* points along each dimension, the number of entries in a transition matrix will be *L*^2(*M*−1)^. For example, for 8 genotypes and a lattice resolution of just 3 points per genotype dimension, the number of elements in a transition matrix is 3^2(8−1)^ = 3^14^≈ 5×10^6^. Of course, this transition matrix is very sparse, containing only ≲ 2(*M*− 1)*L*^*M*−1^ non-zero elements (this upper-bound overcounts for boundary states), which means for 8 genotypes and *L* = 3 there are ≲ 2(8 1)(3)^(8−1)^ = 14(3)^7^ ≈3 ×10^4^ non-zero elements in the transition matrix. Note that the population size *N* never appeared in these estimates; in contrast, for full WF dynamics with population size *N*, the number of non-zero transition matrix elements is the number of multinomial coefficients for *N* objects into *M* genotype groups, which for *N* ≫ *M* is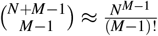, growing exponentially with the size of the population. For *M* = 8 genotypes and a population size of *N* = 10000 (as we use later in the paper), there are 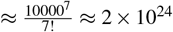non-zero transition matrix elements. These sample calculations demonstrate the significant reduction in computational expenditure required for SHEPHERD compared to trying to calculate optimal policies on full WF dynamics.

### 2.2 Markov Decision Processes for Evolutionary Control

We briefly review the mathematics of Markov Decision Processes (MDPs), which provides a principled approach for optimizing sequential decisions under stochastic dynamics^53^. An MDP is defined by a set of control actions{*a*} exerted on a stochastic system with states {*n}*, which evolves according to Markov transition probabilities *W* (*m*|*n, a*) defining the probability to transition from state *n* to state *m* under action *a*. A decision is made at each time step to perform action *a* with probability *π*(*a*|*n*), conditioned on the current state of the system, which is called the policy.

One often wishes to maximize the expected discounted cumulative reward 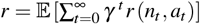where *r*(*n, a*) is the instantaneous reward for state *n* and action *a*, and the discount factor *γ∈* [0, 1) quantifies the desired time-horizon (*γ* = 0 maximizes the instantaneous reward, whereas the cumulative infinite-time rewards are maximized in the limit as *γ* approaches 1). The optimal policy *π*^*\**^(*a*| *n*) yielding this maximized reward is actually deterministic^53^, i.e., it becomes a function *a*^*\**^(*n*) dictating the single optimal action *a*^*\**^ for a given state *n*.

In our context, the state space consists of genetic compositions of the disease (by which we mean, more accurately, the discretized and coordinate-transformed genotype–frequency distributions ***u***^(*n*)^ discussed above in Section 2.2). Choosing drug *a* in state *n* determines the transition probability matrix *W* ^(*a*)^ for that time step Δ*t*, encoding the stochastic evolutionary dynamics under the selective pressure of that drug, as depicted schematically in **Figure 1**. To capture the therapeutic goal of minimizing disease population fitness, we define the instantaneous reward for our MDP as the negative mean fitness of the disease population,

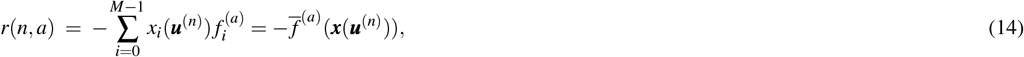

where *x*_*i*_(***u***^(*n*)^) is the frequency of genotype *i* at lattice site *n*, using the inverse of the coordinate transformation discussed above, and 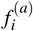is its reproductive fitness under drug *a*. With this definition, higher-fitness populations yield lower (more negative) rewards, so that maximizing the cumulative reward is equivalent to suppressing evolutionary adaptation. We adopt the infinite-horizon discounted formulation, in which future rewards are weighted by a discount factor *γ* = 0.99 (set slightly less than 1 for numerical tractability). This setting emphasizes long-term evolutionary control while ensuring the optimization problem is well-posed and calculable. The optimal MDP policy can then computed numerically, using the well known value iteration algorithm^53^, yielding the optimal drug to administer as a function of genetic composition in order to minimize long-term average fitness of the disease populations. Note that this policy depends only on the genetic composition at that time and not on time itself, thus it is called a stationary policy.

### 2.3 Closed-Loop Implementation in Wright–Fisher Simulations

To evaluate the effectiveness of our approximations in computing the MDP-derived control strategy that we have termed SHEPHERD, we implemented the resulting optimal policy in simulations of the exact WF evolutionary dynamics. While the optimal MDP-derived strategy is mathematically guaranteed to beat any other drug policy under our approximations, the implementation on exact WF simulations provides a nontrivial validation of our approximations.

We begin with three example systems increasing in number of genotypes; for each drug *a*, genotype-specific fitness values 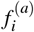were uniformly randomly generated within the very-weak-selection regime 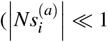for all genotypes *i* and drugs *a*, where the selection coefficients are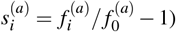, ensuring small selective differences and maintaining population heterogeneity; recall that this is in contrast to the strong-selection, weak-mutation (SSWM) regime employed by previous work^58^, in which the opposite limit is assumed of 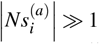 for all genotypes and drugs, and new mutations fix in a population so quickly as to make the population effectively perpetually homogeneous. Furthermore, the fitness landscapes used here incorporate drug-dependent trade-offs such that no single drug minimizes fitness across all genotypes, making the evolutionary control problem inherently non-trivial (see Supplementary Information IV for exact fitness values for each example setup). In other words, these synthetic fitness landscapes exhibit collateral sensitivity between drugs, allowing for the ability to nontrivially steer the evolution of the disease. Recall that WF dynamics tends toward increasing population fitness, so it is only with collaterally sensitive drugs where the fitness ranking of genotypes flips between drugs that we can effectively trap a disease population in a lower fitness state^76^.

Unless otherwise noted, all simulations were initialized with the entire population at the wild-type genotypes (this loses no generality, since absorbing states would not occur with these dynamics, thus long-term behavior does not depend on initial condition). At each generation, the current genotype-frequency vector ***x***_*t*_ was mapped to the nearest coordinate-transformed, discretized state ***u***^(*n*)^. The MDP policy *a*\*(*n*) was then consulted to determine the optimal drug to administer at that state. The chosen drug defined the fitness landscape for the subsequent WF update, which proceeded through the standard steps of selection, mutation, and multinomial sampling with population size *N*. This procedure was iterated for successive generations, thereby creating a closed-loop control system in which therapeutic decisions adapt dynamically to the evolving population state (**Figure 1D** ).

For comparison, we also simulated fixed-treatment strategies in which a single drug was applied continuously, as well as alternating two-drug switching policies. In the latter case, the drugs alternated every generation (e.g., generation *t*: drug A, generation *t* + 1: drug B, then back to A, and so on). We then analyzed the mean fitness trajectories across replicates (to account for the inherent stochasticity) to assess how the MDP-guided control influenced evolutionary dynamics relative to both constant single-drug therapy and alternating two-drug therapy. Because the two-drug policy produces oscillatory fluctuations in fitness, we compared the SHEPHERD trajectories against the two-generation average fitness of the switching strategy.

## 3 Results

### 3.1 Optimal Control in Small Genotype Spaces

We begin with a simple heterogeneous system containing *M* = 3 genotypes connected through symmetric mutation (see Supplementary Information I). While such a system might seem less biologically realizable, it nevertheless provides a strong initial demonstration of our methods, after which we will explore examples with potentially greater biological relevance.

The optimal MDP policy depends explicitly on the current genotype frequency, with different regions of the genotype simplex assigned to different drugs (**Figure 2A,B**). Panel A shows the genotype-frequency space colored by the drug selected by the MDP, with arrows indicating the deterministic part of the evolutionary dynamics under the optimal MDP policy. Rather than driving the population to fixation at a resistant corner, the dynamics converge toward a recurrent region in the center of the simplex, where the stationary distribution is localized (**Figure 2B**); note that this stationary distribution represents highly genetically heterogeneous disease – exactly as we set out to model – as it resides near the center of the simplex, rather than near one of the corners (single-drug policies do, in fact, steer the evolution towards corners representing the genotypes most resistant to those drugs).

**Figure 2.**
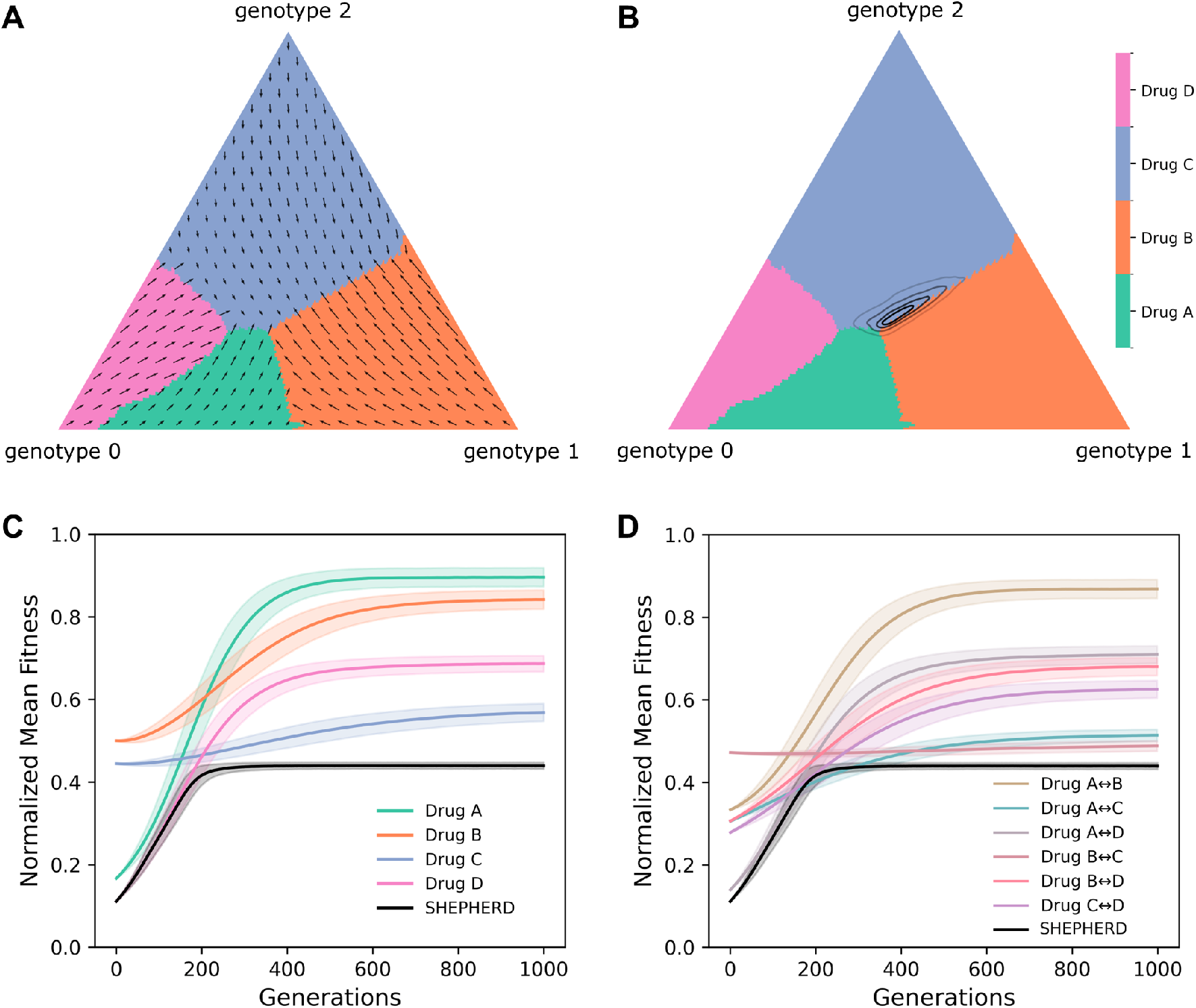
MDP-guided control in a three-genotype system with symmetric mutation. **A**. Optimal drug policy on a synthetic fitness landscape with three genotypes and four drugs, assuming equal mutation rates between all genotypes. Each region of the genotype-frequency simplex (*x*_0_ + *x*_1_ + *x*_2_ = 1) is colored by the choice of drug to minimize long-term mean fitness. Arrows indicate the deterministic direction of population evolution under the corresponding drug. **B**. Long-term stationary distribution of the population under the optimal MDP policy, showing where the evolutionary dynamics concentrate in the long-term limit. Contour lines denote probability density levels of 0.02, 0.20, 0.50, and 0.80. **C**. Wright–Fisher simulations (10,000 replicates) comparing mean fitness trajectories under the optimal MDP strategy versus fixed single-drug treatments. **D**. Wright–Fisher simulations (10,000 replicates) comparing the MDP policy to alternating two-drug switching strategies (e.g., A–B–A–B). In panels C and D, solid lines denote the mean normalized fitness and shaded regions represent standard deviations across replicates. Normalized mean fitness is scaled such that the minimum fitness across all available drugs maps to 0 and the maximum fitness maps to 1. The MDP-guided policy maintains substantially lower long-term fitness than any fixed or switching drug regimen.

We then compared the adaptive control strategy against constant single–drug treatments by analyzing normalized mean fitness over time (**Figure 2C**). Here, normalized mean fitness rescales the population’s average reproductive fitness to the range [0, 1] across all available drugs. This quantity is directly related to classical genetic load^77^, which is defined as the reduction in mean population fitness relative to the maximum attainable fitness. Thus, lower normalized fitness corresponds to higher genetic load. The MDP-guided policy maintains the population at consistently lower fitness levels than any single-drug regimen, achieving long-term suppression of resistance. We also benchmarked the optimal MDP against alternating two-drug switching policies of the form A–B–A–B (**Figure 2D**). Although two-drug switching policies can sometimes achieve lower fitness than single-drug therapies, their fitness trajectories exhibit oscillatory fluctuations, and their long-term averages remain higher than the MDP-guided strategy. Thus, the adaptive control policy not only outperforms fixed therapies but also achieves lower long-term fitness than any periodic switching schedule.

### 3.2 Evolutionary Control with Structured Mutation Networks

We next extended our analysis to a four-genotype fitness landscape connected by Hamming-1 mutation (**Figure 3A**). In this landscape, each genotype can mutate only to its single-bit neighbors (i.e. only mutations in a single gene occur per generation, see Supplementary Information I), resulting in more constrained connectivity compared to the fully symmetric three-genotype case. This four-genotype system thus represents a combinatorially complete set of genotypes with two possible mutations that are either present or absent. To visualize the optimal MDP policy, we sliced the 3D genotype-frequency simplex (*x*_0_ + *x*_1_ + *x*_2_ + *x*_3_ = 1) along the *x*_3_ axis, producing a series of triangular cross-sections. Each slice is colored according to the drug predicted by the MDP to minimize long-term mean fitness, revealing nontrivial policy boundaries that shift as the abundance of genotype 3 varies.

**Figure 3.**
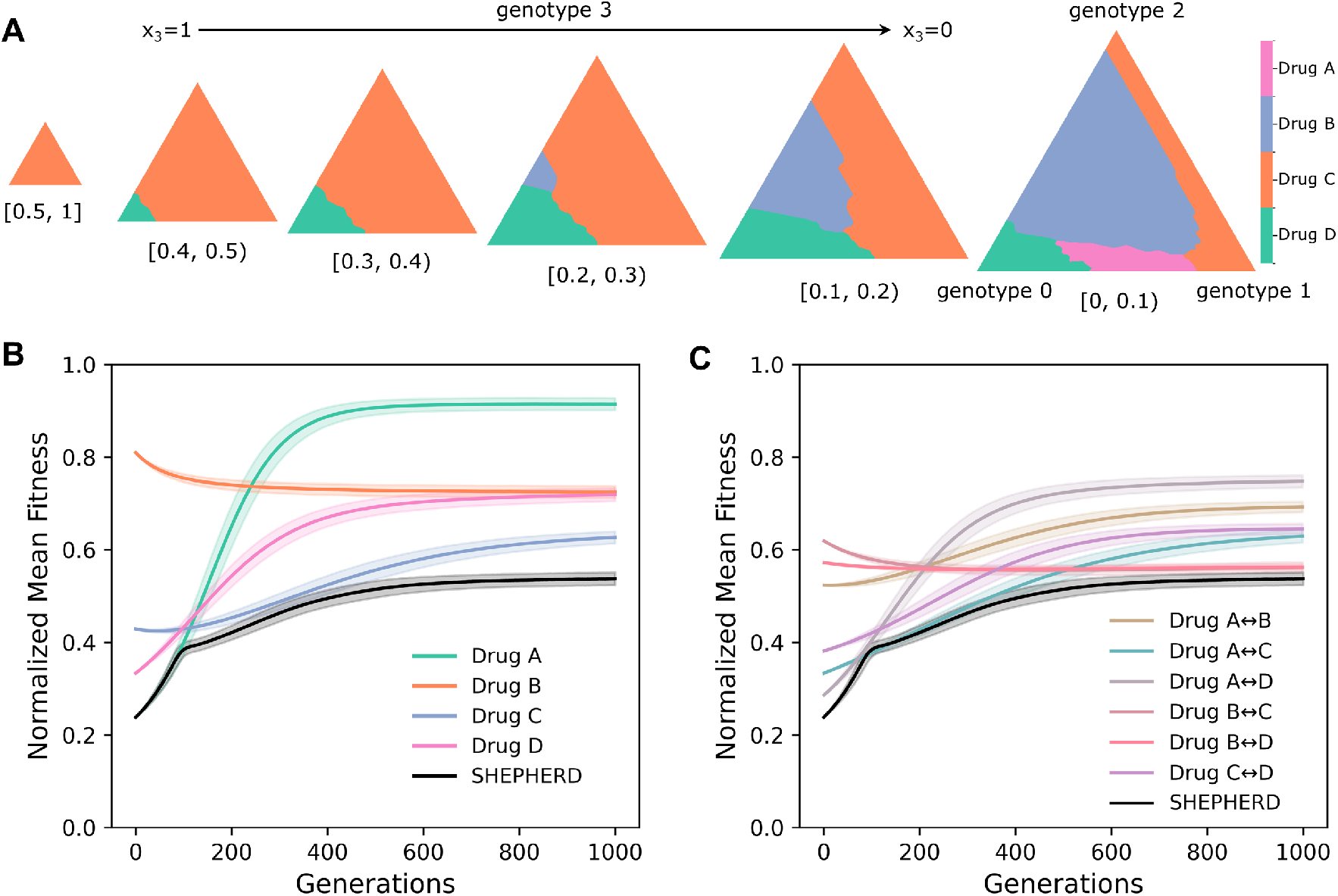
MDP-guided control in a four-genotype system with Hamming-1 mutation. **A**. Optimal drug policy on a synthetic four-drug, four-genotype fitness landscape, assuming mutations occur only between genotypes at Hamming distance 1. The 3D simplex (*x*_0_ + *x*_1_ + *x*_2_ + *x*_3_ = 1) is visualized by slicing along the *x*_3_ axis from *x*_3_ = 1 (left) to *x*_3_ = 0 (right). Each triangular slice is colored by the drug chosen to minimize long-term mean fitness at that genotype composition. **B**. Wright–Fisher simulations (10,000 replicates) comparing normalized mean fitness trajectories under the optimal MDP strategy versus constant single-drug treatments. **C**. Wright–Fisher simulations (10,000 replicates) comparing the MDP policy to alternating two-drug switching strategies (e.g., A–B–A–B). In panels B and C, solid lines denote the mean normalized fitness and shaded regions represent standard deviations across replicates. Normalized mean fitness is scaled such that the minimum fitness across all available drugs maps to 0 and the maximum fitness maps to 1. The MDP-guided policy consistently maintains lower long-term fitness than any fixed or switching drug regimen.

We then evaluated the performance of the MDP-guided strategy in WF simulations. Compared to fixed single-drug treatments, the optimal MDP maintains consistently lower normalized mean fitness across replicates (**Figure 3B**), demonstrating that adaptive control continues to suppress resistance more effectively even as the genotype space grows in complexity. We also compared the MDP to alternating two-drug switching policies (**Figure 3C**). Although some switching strategies transiently reduce fitness below that of constant single-drug therapies, their trajectories exhibit oscillatory fluctuations and their long-term averages remain higher than the optimal MDP. Thus, in this four-genotype landscape, the adaptive policy again achieves the lowest long-term fitness, outperforming both fixed and switching treatments.

These results demonstrate that the MDP framework generalizes beyond the minimal three-genotype case: when applied to a standard four-genotype fitness landscape with Hamming-1 mutation network, adaptive control maintains its advantage by dynamically responding to the evolving population state.

### 3.3 Performance in Higher-Dimensional Fitness Landscapes

Finally, we analyzed an eight-genotype system with Hamming-1 mutation (see Supplementary Information I) (**Figure 4**); this system thus represents a combinatorially complete set of genotypes with three possible mutations that are either present or absent. In higher-dimensional fitness landscapes, the number of possible genotype-frequency states increases exponentially as discussed above, so we employed coarser lattice discretizations to keep the transition matrices computationally tractable. This reduced resolution modestly diminishes the performance of the MDP-guided strategy compared to lower-dimensional cases.

**Figure 4.**
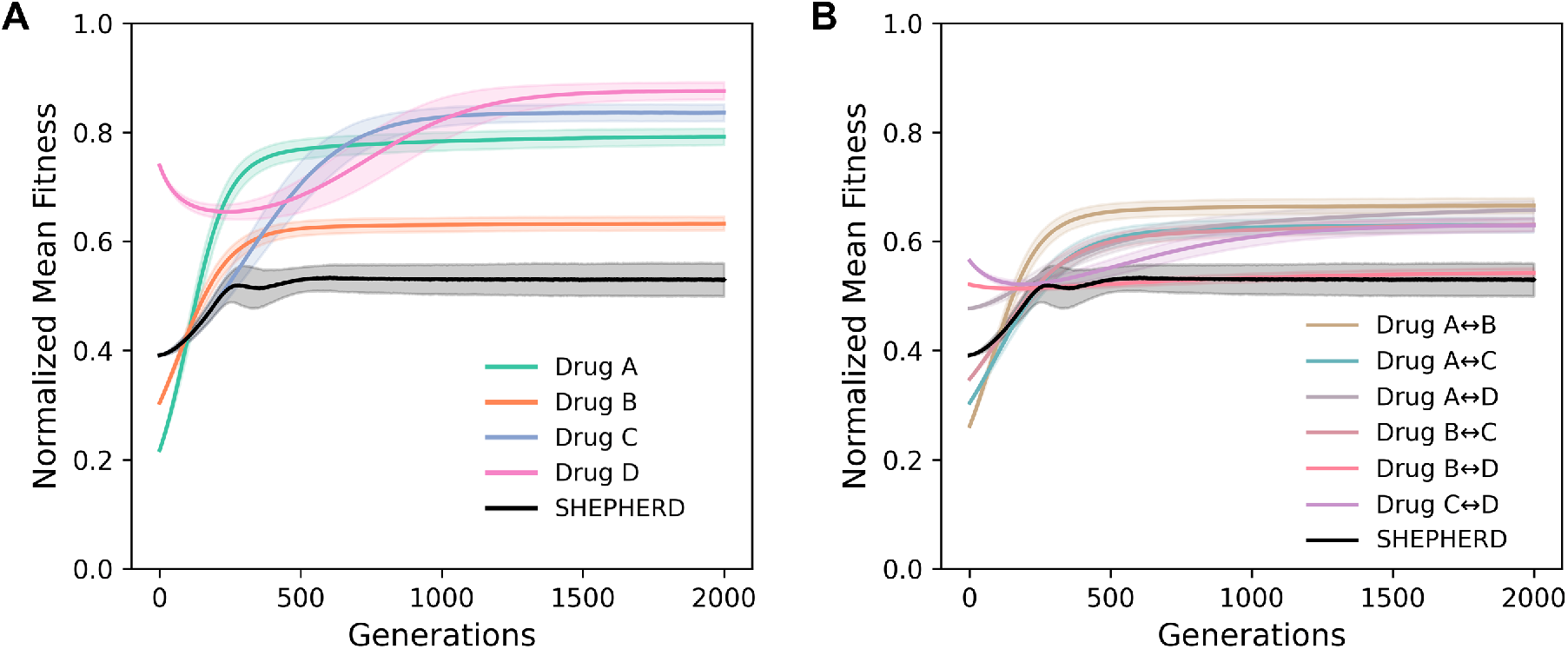
MDP-guided control in an eight-genotype system with Hamming-1 mutation. **A**. Wright–Fisher simulations (10,000 replicates) comparing normalized mean fitness trajectories under the optimal MDP strategy versus fixed single-drug treatments in a synthetic fitness landscape with eight genotypes and four drugs, assuming mutations occur only between genotypes at Hamming distance 1. **B**. Wright–Fisher simulations (10,000 replicates) comparing the MDP policy to alternating two-drug switching strategies (e.g., A–B–A–B). In both panels, solid lines denote normalized mean fitness and shaded regions represent standard deviations across replicates. Normalized mean fitness is scaled such that the minimum fitness across all available drugs maps to 0 and the maximum fitness maps to 1. The MDP-guided policy maintains lower long-term fitness than any constant single-drug or alternating two-drug strategy.

Even so, Wright–Fisher simulations show that the optimal MDP policy continues to outperform both fixed single-drug regimens and alternating two-drug switching strategies (**Figure 4**). Although the margin between adaptive control and the best switching strategies narrows in this more complex setting, the MDP consistently achieves the lowest long-term mean fitness across replicates. Interestingly, for both Drug D monotherapy and the C ↔ D alternating regimen, we observed an initial dip in fitness followed by a rise toward the global maximum. This behavior reflects a transient valley crossing, in which populations temporarily pass through lower-fitness intermediates before reaching a resistant peak – a phenomenon known as stochastic tunneling^66,78^.

These results underscore both the scalability and the limitations of our approach: while discretization becomes more challenging in large genotype spaces, adaptive evolutionary control remains more effective than heuristic fixed or switching therapies.

### 3.4 Sensitivity and Robustness of SHEPHERD

To assess how modeling choices impact performance, we quantified the sensitivity of the MDP-guided control to (i) the lattice discretization used to construct the transition matrices (**Figure 5A**) and (ii) the temporal resolution of therapeutic decisions, quantified by the drug update interval Δ*t* (**Figure 5B**).

**Figure 5.**
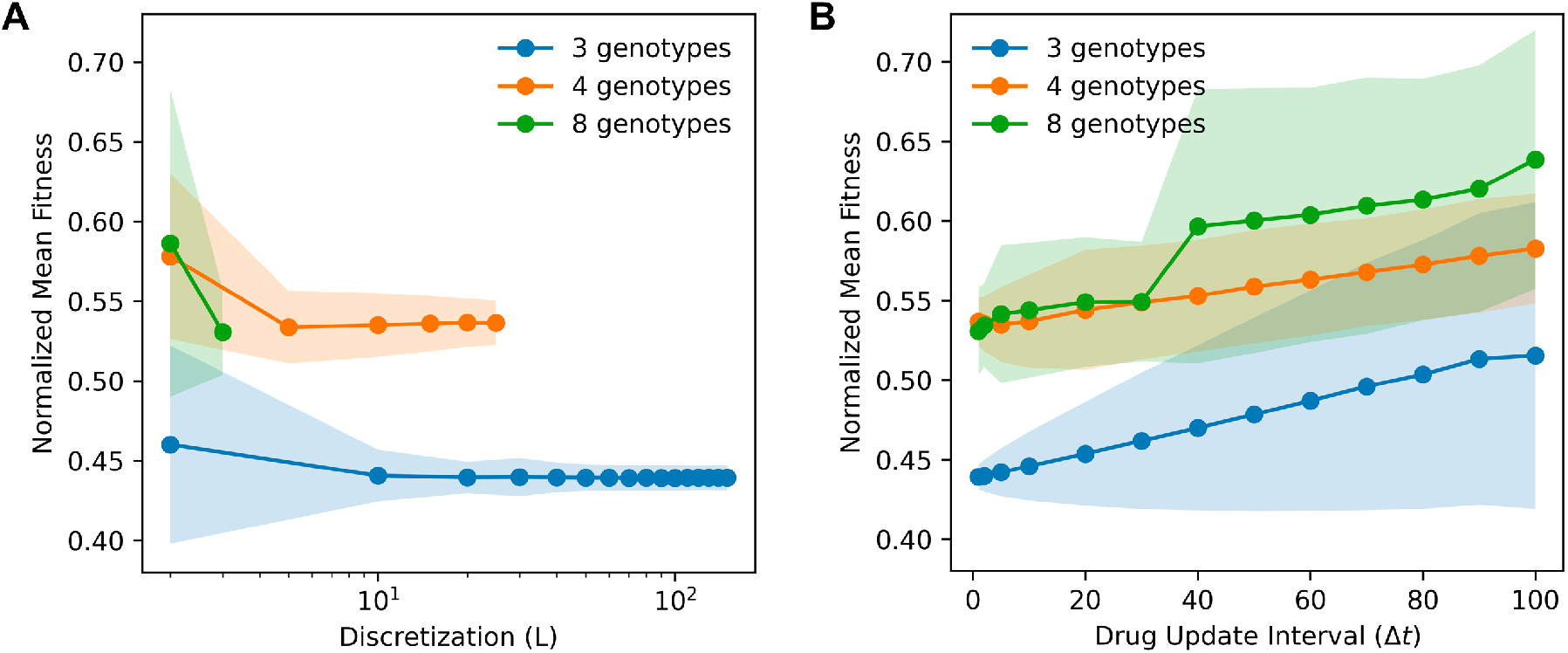
Sensitivity and Robustness of SHEPHERD to spatial and temporal resolution. **A**. Effect of lattice discretization (*L*) on long-term normalized mean fitness for systems with 3, 4, and 8 genotypes, using Δ*t* = 1. Increasing *L* (finer discretization) improves the fidelity of the discretized dynamics, lowering fitness until performance plateaus at finite resolution. **B**. Effect of the drug update interval (Δ*t*; Wright–Fisher generations per decision) on normalized mean fitness, with discretization set to *L* = 100, 20, and 3 for the 3-, 4-, and 8-genotype systems, respectively. Larger Δ*t* (less frequent therapeutic updates) degrades performance in all genotype spaces. In both panels, solid lines denote means across replicates and shaded regions show standard deviations. Normalized mean fitness is scaled within each experiment so that the minimum and maximum across available drugs map to 0 and 1, respectively.

#### Discretization (L)

Increasing the lattice resolution improves the fidelity of the discretized Fokker–Planck dynamics and yields lower long-term normalized mean fitness across genotype spaces (**Figure 5A**). In the three- and four-genotype systems, performance rapidly converges as *L* increases, and the variance across replicates becomes progressively smaller, indicating that finite discretizations are sufficient for stable policies. For the eight-genotype system, we were limited to coarser grids (*L* = 2, 3) due to computational constraints, but the same qualitative trend is evident: increasing resolution lowers fitness and reduces variability, although performance seems to plateau once a finite resolution is reached.

#### Drug update interval (Δt)

Coarser temporal control degrades performance in all genotype spaces (**Figure 5B**). As Δ*t* increases, the controller has fewer opportunities to adapt to the evolving composition, leading to progressively higher normalized mean fitness and wider variability across replicates. The effect is most pronounced in the eight-genotype landscape, reflecting the faster movement through a larger state space under fixed drugs when updates are infrequent. Previous work exemplified similar results regarding evolutionary control under imperfect monitoring^30^, although that work considered only two genotypes and two control actions.

Taken together, these analyses highlight two practical guidelines: (i) use a lattice fine enough to achieve policy convergence in the target dimension, and (ii) make drug decisions as frequently as is clinically feasible (small Δ*t*), since rapid feedback most effectively suppresses adaptation. These results corroborate the intuition that control can be exerted more effectively when the controller has better information, as afforded by more frequent or more resolved observations of the genetic composition of a disease; this is a fruitful topic for further study.

### 3.5 Broad Applicability of SHEPHERD

The strength of mutation in evolutionary dynamics is quantified by the product *Nµ*, where *N* is the population size and *µ* is the per-generation mutation rate (which we assume here to be constant across all accessible mutations): we consider *Nµ* < 1 the weak mutation regime, and contrastingly *Nµ* > 1 defines the strong mutation regime. Similarly, the strength of selection is quantified by the products 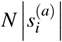, where 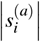is the magnitude of the selection coefficient for genotype *i* under drug *a*: when 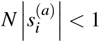for all genotypes and drugs, we consider that weak selection, whereas 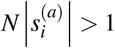for any genotype and drug puts us in the strong selection regime. (Note that the definitions of weak and strong mutation and selection are looser in this section, with less extreme inequalities to cover broader ranges of evolutionary conditions.)

We initially set out specifically to compute optimal drug protocols for controlling disease populations beyond the extremes of the strong-selection-weak-mutation (SSWM) regime, which had been explored previously^58^, as we discussed above. Here we wish to emphasize the broad applicability of the approach we developed.

We tested SHEPHERD under a wide variety of evolutionary regimes using synthetic fitness landscapes defined on four genotypes and four drugs. Specifically, we considered all four combinations of weak or strong selection and weak or strong mutation. For four drugs and four genotypes, we generated fitness landscape sets by randomly drawing fitness values normally distributed around 1 for each of the 16 genotype/drug pairs, then sorted the resulting landscape set into weak or strong selection according to the criteria given at the beginning of this subsection (see Supplementary Information IV); we used this procedure to generated 100 viable fitness landscape sets in each mutation/selection regime (400 total), explicitly excluding landscape sets containing a single drug effective against all genotypes (because the optimal policy trivially employs only that single drug), and discarding fitness values below 0. Evolutionary dynamics on each landscape set were simulated under both weak and strong mutation using WF dynamics with population size *N* = 10000, and long-term mean fitness was evaluated from 1000 simulation replicates per landscape set in the last 100 generations of a 1000 generation simulation. Across all regimes considered, SHEPHERD consistently achieves equal or improved long-term performance relative to the best fixed two-drug policy, as summarized in **Figure 6**. Note that the example discussed in Section 3.2 and depicted in **Figure 3** would be merely one point in the histogram of **Figure 6C**.

**Figure 6.**
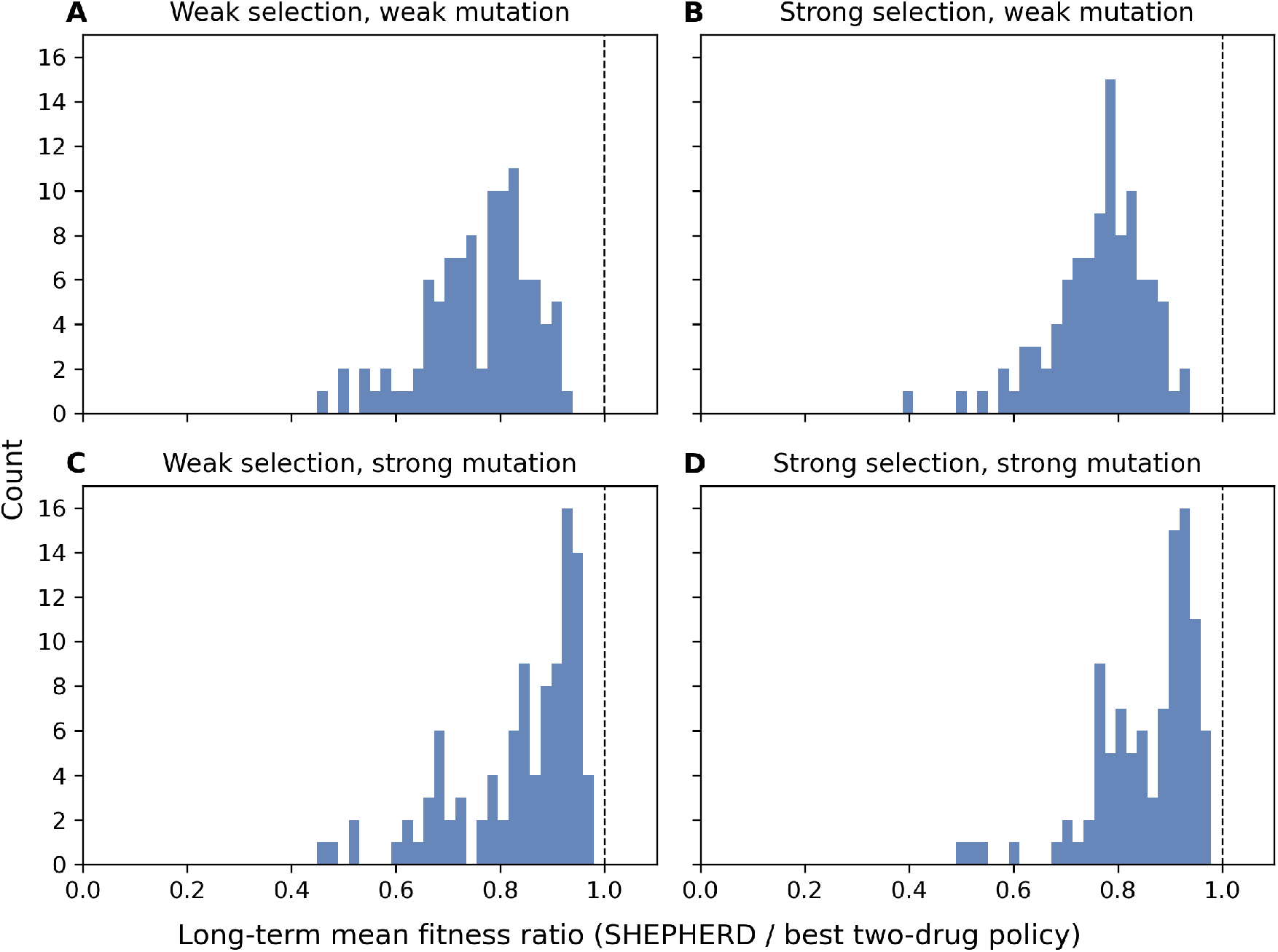
SHEPHERD is successful across selection and mutation regimes. Histograms show the distribution of the long-term mean fitness ratio (SHEPHERD / best two-drug policy) across four evolutionary regimes defined by weak or strong selection and weak or strong mutation (A–D). Each panel summarizes results from 100 independently generated fitness-landscape sets, each consisting of four drugs acting on four genotypes. For each landscape set, evolutionary dynamics were simulated using 1000 Wright-Fisher simulation replicates, each with population size *N* = 10,000. Weak mutation regimes correspond to a mutation rate *µ* = 5 × 0^−5^ (*Nµ* < 1), while strong mutation regimes correspond to *µ* = 10^−3^ (*Nµ* > 1). the magnitude of selection coefficients: weak selection corresponds to 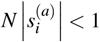for all genotypes *i* and drugs *a*; otherwise, Long-term mean fitness was computed by time-averaging over generations [9901, 10000]. Selection regimes were defined by the system is in the strong selection regime. Fitness landscape sets were generated by randomly drawing fitness values normally distributed around 1 for each of the 16 genotype/drug pairs, then sorting the resulting landscape set into weak or strong selection (see Supplementary Information IV). For the fitness ratio depicted on the horizontal axis of these histograms, values less than or equal to 1 indicate successful control by SHEPHERD. Across all regimes, SHEPHERD matches or outperforms the best two-drug policy.

This analysis encompassed in **Figure 6** suggests the potential power of SHEPHERD beyond its initial motivations. Many systems across the spectrum of biology obey the Wright-Fisher dynamics—or more precisely the Fokker-Planck dynamics obtained in the large population limit of Wright-Fisher—underlying SHEPHERD. The mutational dynamics may differ, but SHEPHERD is amenable to varied mutational topologies: the three-genotype example took place on a fully connected mutational graph, in which any genotype could mutate to any other genotype in one time-step, whereas the four- and eight-genotype examples took place on hypercubic mutational graphs, in which genotypes only mutate to their direct mutational neighbors in one time-step. Additionally, the objective function to be optimized by SHEPHERD can be arbitrarily defined, allowing for alternate or complementary goals such as promoting polymorphism^30^ or minimizing the toxicity and side effects of therapeutic interventions^79^. Relatedly, while we spoke of switching between drugs in our examples, another dimension of control is the continuous modulation of drug doses; our general formulation of control actions is immediately amenable to describing the same drug administered at distinct doses, or even fitness under the lack of any drug. Plus, therapeutic interventions like drugs and radiation are merely one of many possible external stimuli with the power to steer evolution (e.g. temperature, presence of nutrients or metabolites), any of which are amenable to optimization with SHEPHERD. Our approach is thus generalizable beyond controlling evolving diseases with drug protocols—it could compute optimal control strategies for a wide class of biological phenomena^5^, and even go beyond biology to inform evolutionary computation^80^. Further uses of SHEPHERD in varied situations is another fruitful topic for future study, including studying which aspects of landscape sets translate to higher fitness reductions under optimal protocols^76^.

## 4 Discussion

In this work, we develop SHEPHERD, a framework for adaptive evolutionary control that integrates the stochastic Wright–Fisher (WF) model of heterogeneous populations with the optimization power of Markov Decision Processes (MDPs). By coupling the diffusion approximation of WF with a useful coordinate transformation and subsequent lattice discretization, we efficiently derived optimal control strategies under the MDP framework—in our examples, drug policies that minimize long-term mean disease fitness across different genotype spaces with varied fitness landscapes and different mutational dynamics. While the optimality of these policies is guaranteed by definition when calculated exactly, WF simulations confirmed that our adaptive strategies calculated under approximation still outperform constant single-drug regimens and alternating two-drug schedules, even in higher-dimensional fitness landscapes where resistance pathways are more complex^81^. These results demonstrate that adaptive evolutionary control can be systematically designed and evaluated in genetically heterogeneous settings.

Furthermore, due to the mathematical optimality of MDP-derived policies, these results should hold in general, for any underlying fitness landscapes: the optimal MDP-derived policy will always perform at least as well as a single-drug policy by definition, as long as the underlying approximations are valid. There is one situation, however, in which an optimal adaptive policy would not outperform a single-drug policy—when the optimal adaptive policy *is* a single-drug policy. This case would correspond to the most fit genotype under that single drug still being less fit then genotypes that have lower fitness under all other drugs; this case is, in some sense, trivial. Thus we have explicitly chosen underlying fitness landscapes exhibiting collateral sensitivity, in which genotypes have opposite fitness rank under different drugs; this allows for evolutionary steerability, providing for therapeutic benefit of adaptively switching between drugs.

A central contribution of SHEPHERD is that it relaxes the *strong-selection weak-mutation (SSWM)* assumption, under which populations are effectively homogeneous and evolve through successive fixations. Prior MDP-based control strategies have operated in this simplified setting^23,58,76^. By contrast, the general WF framework allows the coexistence of multiple genotypes, with mutations segregating concurrently, providing a more realistic model of genetic heterogeneity. In such heterogeneous settings, SHEPHERD is particularly effective at capturing this diversity and suppressing the evolution of resistance by reducing long-term fitness. When a system truly resides in the SSWM limit, accurate modeling might need extremely fine lattice discretization near simplex corners to represent nearly homogeneous states, reducing efficiency. In such cases, SHEPHERD could be slower to compute, and approaches explicitly tailored to SSWM dynamics^23,58,76^may be more appropriate. However, as we demonstrated by testing SHEPHERD across a multitude of conditions, it is expected to perform successfully under broad evolutionary regimes.

From a computational perspective, discretization remains the major bottleneck of our method. The number of states grows rapidly with both genotype dimension and lattice resolution, rendering value iteration increasingly expensive. For higher-dimensional landscapes, alternative methods such as reinforcement learning could provide a promising path forward, enabling approximation of optimal policies without exhaustive discretization. Such approaches may balance tractability and performance, particularly when scaling to realistic genotype spaces relevant to clinical applications. In addition to added dimensionality, future work could incorporate interactions in the form of frequency-dependent fitness, which is known to confound evolutionary expectations^67,82^, as well as a continuous action-space to model the important effects of continuously varying drug dosages^83^.

Finally, our current results rely on synthetic fitness landscapes, as empirical measurements of genotype-specific fitness under multiple drugs in non-SSWM regimes are, to our knowledge, not yet available. Experimental efforts to map such landscapes would provide critical data for validating and refining our framework. Despite this limitation, our study offers a conceptual advance by showing how adaptive control can be extended from homogeneous, SSWM models to broader evolutionary settings, highlighting both the opportunities and the challenges in designing optimal evolutionary therapies, and generally in controlling evolving systems.

## Acknowledgments

Research reported in this publication was supported by the National Cancer Institute of the National Institutes of Health under Award Number T32CA094186 (J.A.P.), R37CA244613 (J.G.S.), and by the American Cancer Society under Grant RSG-20-096-01 (J.G.S.). The content is solely the responsibility of the authors and does not necessarily represent the official views of the National Institutes of Health. The drug icons used in Figure 1D were created with BioRender.com.

## Supplementary Information

### I. Mutation Matrices for Example Setups

The structures of 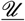used in this study, along with their corresponding mutation graphs, are shown in Figure SI 1.

**Figure SI.1.**
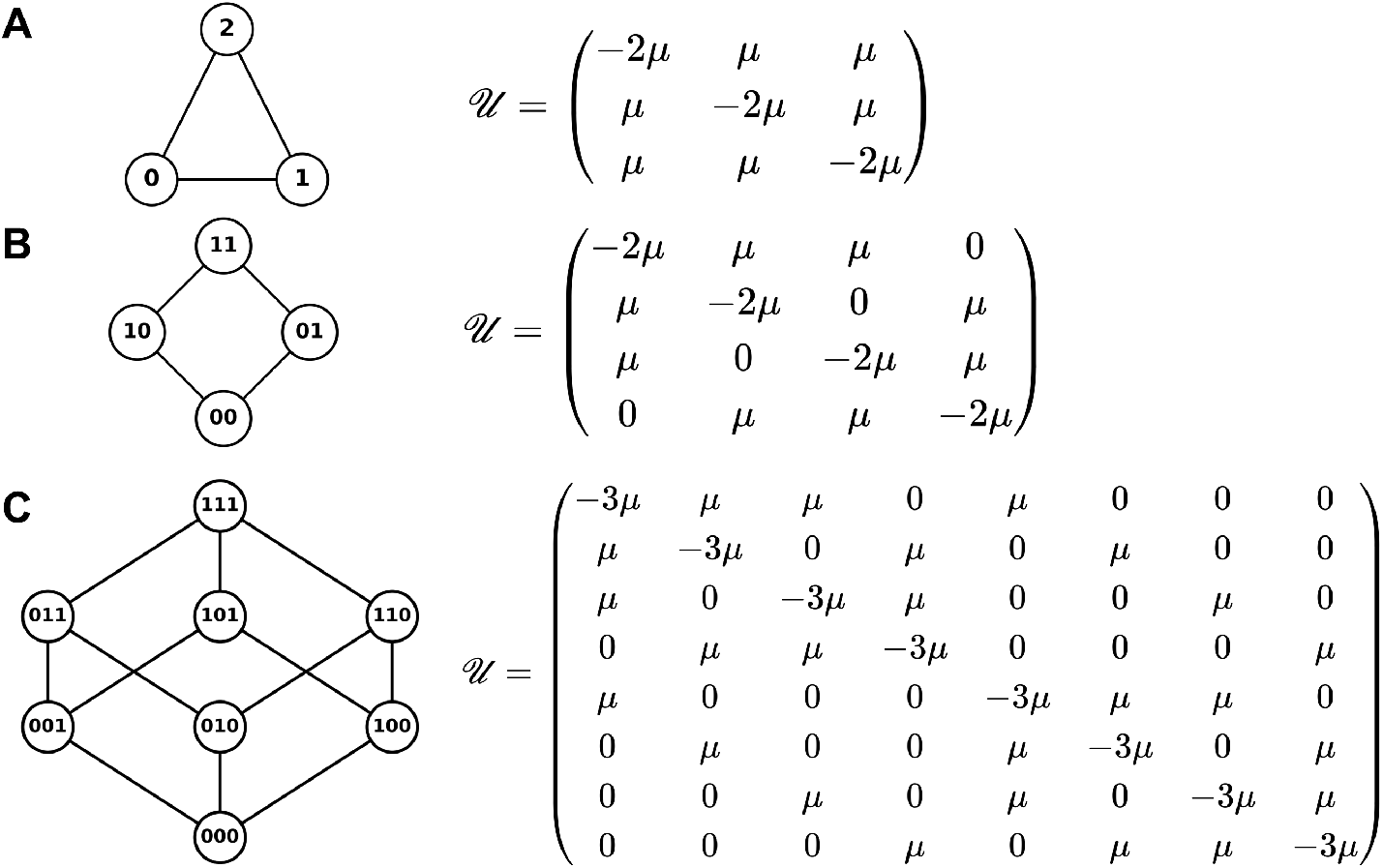
Mutation graphs and corresponding matrices 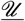for (A) three-, (B) four-, and (C) eight-genotype systems used in this study. Each node represents a genotype, and connected edges indicate possible mutations occurring symmetrically at rate *µ*. The corresponding mutation matrices 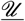are shown to the right, with diagonal entries *U*_*ii*_ ensuring each column sums to zero.

#### II. Derivation of Deterministic Part of Fokker-Planck Equation in Transformed Coordinates

In the new coordinates, the deterministic part, like the diffusion part, follows from the Itô coordinate transformation of the Fokker–Planck equation^84^. Writing 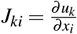(the inverse Jacobian), and noting that the second term arising from noise vanishes by symmetry, we have

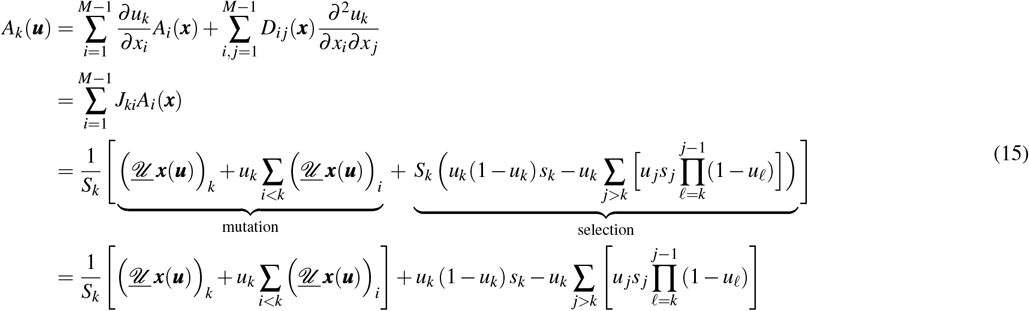

#### III. Derivation of Lattice Discretized Transtion Rates

To connect the continuous Fokker-Planck equation to the lattice formulation used in numerical simulations, we consider a finite lattice spacing *h* along each coordinate *u*_*i*_ and approximate the probability fluxes between nearest-neighbor sites. Let 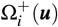and 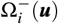denote the forward and backward transition rates along the *i*th direction, respectively. The local master equation for probability flow is

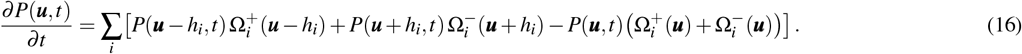

To show that this discrete master equation converges to the continuous Fokker-Planck dynamics, we perform a Taylor expansion of the probability and rate terms for small *h*:

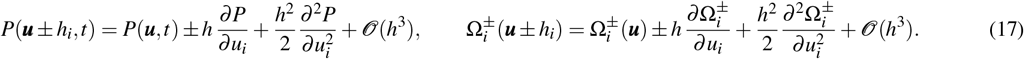

Substituting these expansions into the master equation and collecting terms up to *O*(*h*^2^) yields first-order terms corresponding to the deterministic part and second-order terms corresponding to diffusion.

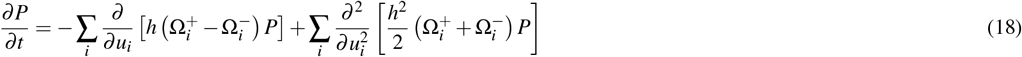

Comparing with the continuous Fokker-Planck equation (6), we derive the correspondence between the deterministic part, the diffusion, and the transition rates:

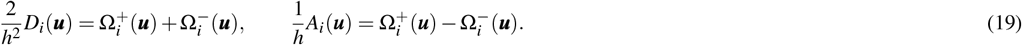

Inverting these yields

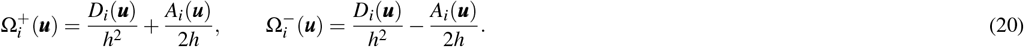

These expressions define the discrete forward and backward transition rates that recover the continuous Fokker-Planck dynamics in the limit *h →* 0, yielding the finite-difference matrix elements discussed in the main text.

### IV. Simulation Parameters and Code Availability

For the three-genotype example, we used a population size of *N* = 5000 with a mutation rate of *µ* = 0.001. For the four- and eight-genotype examples, simulations were performed with population size *N* = 10,000 and the same mutation rate *µ* = 0.001. Each of four drugs defines a distinct fitness landscape, denoted as Drug A–D. Within each landscape, genotype-specific fitness values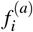 were randomly generated within the stated selection regime (very-weak-selection for the primary examples, in which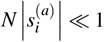 for all genotypes *i* and drugs *a*, i.e. small relative selective differences among genotypes). These synthetic landscapes incorporate drug-dependent trade-offs that ensure no single drug universally minimizes fitness across all genotypes, making the optimal control problem non-trivial. The genotype-specific fitnesses *f* ^(*a*)^ are summarized below for the studied examples with three, four, and eight genotypes, respectively:

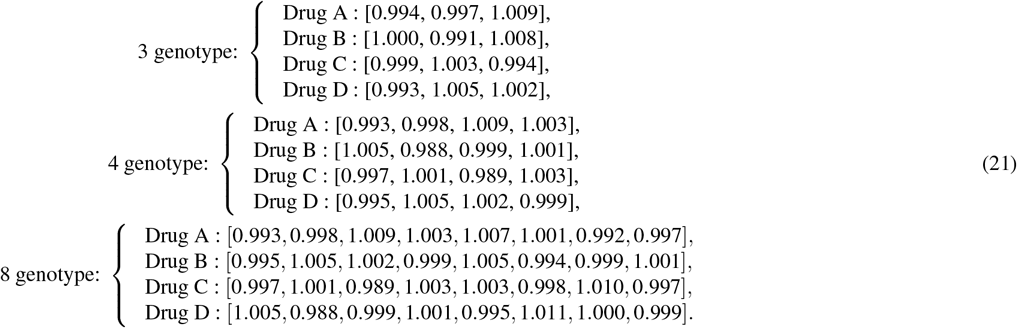

To generate the landscape sets for **Figure 6**, fitness values 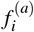for genotype *i* under drug *a* for all 16 genotype/drug pairs comprising four genotypes under four drugs were drawn from normal distributions with mean value 1 and standard deviation 4*/N*, where *N* = 10000 was the population size used for these simulations (fitness values less than 0 were discarded).

The selection coefficients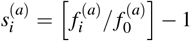were then calculated for each non-wild-type genotype in each drug, the landscape sets were classified as weak-selection if max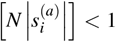, otherwise the landscape sets were classified as and strong-selection. This procedure was used to generate 100 landscape sets for each of the four quadrants of **Figure 6** (400 total landscape sets).

The optimal MDP solution was implemented using value iteration as provided by the Python package `pymdptoolbox` (version 4.0-b3). All code used to generate the results presented in this study is available at https://github.com/Pen9Chen/SHEPHERD.

